# DddY is a bacterial dimethylsulfoniopropionate lyase representing a new cupin enzyme superfamily with unknown primary function

**DOI:** 10.1101/161257

**Authors:** Lei Lei, Uria Alcolombri, Dan S Tawfik

## Abstract

Dimethylsulfide (DMS) is released at rates of >10^7^ tons annually and plays a key role in the oceanic sulfur cycle and ecology. Marine bacteria, algae, and possibly other organisms, release DMS via cleavage of dimethylsulfoniopropionate (DMSP). Different genes encoding proteins with DMSP lyase activity are known belonging to different superfamilies and exhibiting highly variable levels of DMSP lyase activity. DddY shows the highest activity among all reported bacterial lyases yet is poorly characterized. Here, we describe the characterization of recombinant DddY is from different marine bacteria. We found that DddY activity demands a transition metal ion cofactor. DddY also shares two sequence motifs with other bacterial lyases assigned as cupin-like enzymes, DddQ, DddL, DddK, and DddW. These cupin motif residues are essential for DddY activity, as for the other cupin DMSP lyases, and all these enzymes are characterized by a common metal-chelator inhibitor (TPEN). Analysis of all sequences carrying these cupin motifs defined a superfamily: Cupin-DLL (DMSP lyases and lyase-like). The DMSP lyase families are sporadically distributed suggesting that DMSP lyases evolved within this superfamily independently along multiple lineages. However, the specific activity levels, genomic context analysis, and systematic profiling of substrate selectivity as described in the accompanying paper, indicate that for only some of these families, most distinctly DddY and DddL, DMSP lyase is the primary, native activity. In other families, foremost DddQ, DMSP lyase seems to be merely a promiscuous activity. The native function of DddQ, and of nearly all members of this newly identified Cupin-DLL superfamily, remains unknown.

**Abbreviations:**

DMSP: dimethylsulfoniopropionate
DMS: dimethylsulfide
cupin-DLL: cupin DMSP lyase and lyase-like

**Funding:** Financial support by the Estate of Mark Scher, and the Sasson & Marjorie Peress Philanthropic Fund, are gratefully acknowledged. D.S.T. is the Nella and Leon Benoziyo Professor of Biochemistry.

## Introduction

Dimethyl sulfide (DMS) is a key regulator of marine life, and possibly also a climate regulator. DMSP (dimethylsulfoniopropionate) is the primary precursor of DMS, and is synthesized mostly by marine phytoplankton (single-cell algae) but also by macroalgae, corals, some angiosperms and bacteria.^1-5^ DMSP probably acts as osmolyte and antioxidant, although its precise physiological role remains unknown, as is the role of DMS release. ^2, 4^ Also unknown is the relative contribution of different marine species to the global DMS release, or even of bacterial relative to algal mediated release.^6^ Several putative bacterial DMSP lyase families were identified and assigned as ‘*Ddd*’ (DMSP dependence DMS releasing) genes,^7^ as well as one eukaryote family, dubbed Alma, found in organisms such as algae and corals.^8^

We aimed at better understanding of the enzymology and phylogenetics of most abundant bacterial DMSP lyases. The so-far identified *Ddd* gene families include: DddD, a DMSP CoA-transferase-lyase;^7, 9^ DddP that belongs to the M24 proteinase family;^10^ DddQ, DddL, DddW and DddK that have been assigned to the cupin class;^11-14^ and, DddY whose classification remained unknown so far.^4^,^15-17^ The eukaryote Alma DMSP lyase belongs to yet another superfamily (Asp/Glu racemase).^9^ In addition to variable evolutionary origins, these families, and foremost the bacterial enzymes, exhibit highly variable levels of DMSP activity. The reported specific activities seems to range from 0.002 up to 0.028 Units (μM DMS min^-1^ mg^-1^ enzyme) for DddQ ^18-20^ up to 675 Units for DddY.^15, 16^ Kinetic parameters are known for only some of these enzymes, but the specific activities relate to *kcat*/*KM* values that may be as low as or even lower than 1 M^-1^ s^-1^ for DddQ,^18-20^ up to 10^6^ M^-1^ s^-1^ for DddY.^15^,^16^ Indeed, DddY, the most active lyase reported to date,^15, 21^ is likely the least characterized one.

To deepen our understanding the enzymology of DMS release, we have further characterized the DddY DMSP lyase family. In doing so, we found that DddY is evolutionary related to DddQ, DddL, DddW and DddK. Together, these 5 families define a new superfamily of enzymes that share the cupin metal site and jellyroll fold. However, the analyses of the kinetic parameters, the substrate specificity (see accompanying paper),and the genome context, presented here all suggest that while the DddY and DddL represent *bona fide* DMSP lyases, other families, most distinctly DddQ, appear to exhibit DMSP lyase activity as a promiscuous, side-activity. This promiscuous activity stems from shared evolutionary origins and overlapping substrate or/and reaction patterns of members of this newly identified superfamily dubbed the Cupin-DLL (Cupin DMSP lyase and lyase-like) superfamily.

## Results

### Recombinant DddY’s exhibit DMSP lyase activity with *kcat*/*KM* in the range of 10^6^ M^-1^ s^-1^

To better characterize the DddY family, we collected all putative DddY sequences. We used the previously identified *Alcaligenes* DddY**_’_** ^15^,^17^ to search the NCBI database. After filtering (≥ 90% coverage and ≥ 30% amino acids identity) and muscle alignment, a phylogenetic tree was built indicating two major clades that are related although with considerable divergence **(Figure 1A;** 32% and 82% average sequence identity compared to AfDddY). The 1^st^ clade included the previously described *Alcaligenes* and *Desulfovibrio* DddY’s,^15, 16^ while the ^2nd^ clade included primarily *Ferrimonas* and *Shewanella* genes. Two additional sequences were identified that appear as outgroup of the 2^nd^ clade and notably belong to *Synechococcus*, widely spread marine photosynthetic cyanobacteria (31% and 33% sequence identity compared to AfDddY).

**Figure 1.**
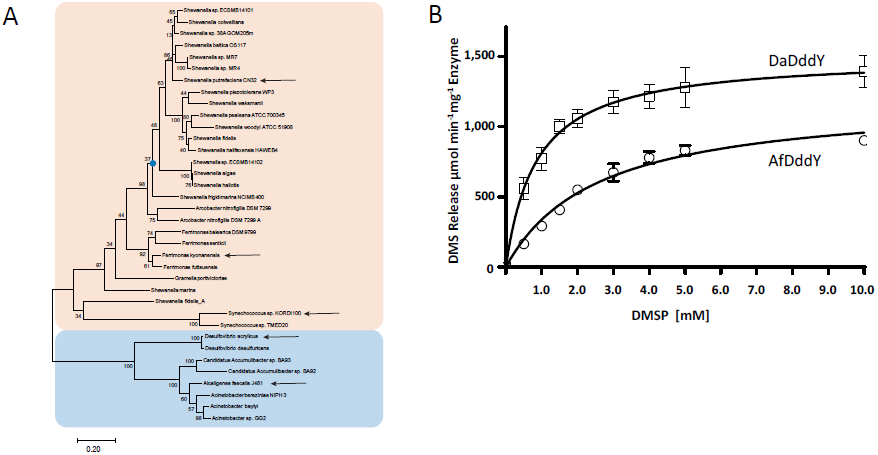
DddY phylogeny and kinetic characterization. (A) A maximum-likelihood tree of identified putative DddY sequences indicates two clades highlighted in pink and blue. Three genes were characterized from the top clade (indicated in arrows), and the ancestral node for the *Shewanella* sub-clade (blue dot) was inferred, yet none of these 4 proteins showed consistent DMSP lyase activity. The bottom clade, however, encompasses highly active DMSP lyases. The Complete gene names and entry numbers are given in **Supplemental dataset 1.** (B) Determination of the kinetic parameters of two recombinant DddY’s from the blue clade: *Desulfovibrio* and *Alcaligenes* DddY’s (DaDddY, AfDddY, respectively). Rates were determined in a buffer containing 1 mM CaCl_2_ under initial rates conditions (see Methods). Data points represent the average specific activity for 3 independent measurements and the error bars represent the S.D. values. Shown is a direct fit to the Michaelis-Menten model (R^2^ ≥ 0.97). The resulting kinetic parameters are presented in **Table 1.**

Representatives of these two clades were cloned and over-expressed in *E. coli*, including *Shewanella putrefaciens* CN-32, *Ferrimonas balearica* DSM 9799, *Desulfovibrio acrylicus* (DaDddY), *Alcaligenes faecalis* J481(AfDddY) and *Synechococcus sp.* KORDI-100 DddY genes. *Alcaligenes* DddY was reported to have a periplasmic signal peptide,^17^ and a homologous N-terminal region seems to appear in all DddY’s. Initially, the wild-type genes including their putative signal peptides were expressed (DNA and amino acid sequences of all constructs are provided as **Supplemental Dataset 1).** Of all tested candidates, DaDddY exhibited the highest activity. However, mass spectrometry of the purified enzyme gave ambiguous results failing to indicate whether the original periplasmic signal peptide was recognized as such in *E. coli*. Thus, the putative mature enzyme sequence was recloned fused to PelB – a widely used *E. coli* periplasmic signal peptide. This recombinant DaDddY variant was purified yielding the 50kDa mature protein at ≥ 95% purity (**Supplemental Fig. S1A).**

Recombinant DaDddY exhibited maximal activity at pH 8.5, although the pH-rate profile indicated multiple titratable groups **(Supplemental Fig. S1B)** The origins of this double-bell-shaped pH-rate profile are unclear, but a very similar profile was reported for endogenous AfDddY.^22^ The recombinant AfDddY variant was obtained with its native signal peptide. Both enzymes obeyed the Michaelis-Menten model **(Figure 1B).** DaDddY exhibited a *kcat*/*KM* value of ~1.3 × 10^6^ M^-1^s^-1^, and its orthologue, AfDddY, showed ~4-fold lower *kcat*/*KM* of 0.35 × 10^6^ M^-1^s^-1^. These values are highly similar to the values originally reported for the enzymes isolated from their endogenous organisms **(Table 1).** Thus, as indicated by previously reported specific activities, DddY shows the highest catalytic efficiency among all known bacterial DMSP lyases. DddY’s catalytic efficiency is also considerably higher than that of Alma algal DMSP lyases (0.8 × 10^5^ M-^1^s-^1^ for *Emiliana huxleyi* Alma1 and 2.7 × 10^4^ M-^1^s-^1^ for *Symbiodinum-*A1 Alma1). ^6, 8^

**Table 1:** Features of the explored DddY and DddL enzymes compared to other cupin-DLL family members

| Family | Species | NCBI Accession | Length (amino acids) | Specific activity ($\mu\text{mol min}^{-1} \text{mg enzyme}^{-1}$) | $k_{cat}/K_M$ ($\text{M}^{-1}\text{s}^{-1}$) | $K_M$ (mM) | $k_{cat}$ ($\text{s}^{-1}$) | Metal cofactor |
| --- | --- | --- | --- | --- | --- | --- | --- | --- |
| DddY | *Alcaligenes faecalis* J481 | E7DDH2.1 | 401 | 902±35 | 3.5*10<sup>5</sup> | 2.56 | 0.9*10<sup>3</sup> | Mn<sup>2+</sup> (2) |
|  |  |  |  | 390 (Ref. <sup>15</sup>) | 2.3*10<sup>5</sup> (Ref. <sup>15</sup>) | 1.41 (Ref. <sup>15</sup>) | 0.3*10<sup>3</sup> Ref. <sup>15</sup> |  |
| | *Desulfovibrio acrylicus* | SHJ73420.1 | 403 | 1391±76 | 1.32*10<sup>6</sup> | 0.85 | 1.13*10<sup>3</sup> | Mn<sup>2+</sup> (2) |
|  |  |  |  | 2100 (Ref. <sup>16</sup>) | 4.60*10<sup>6</sup> (Ref. <sup>16</sup>) | 0.45 (Ref. <sup>16</sup>) | 2.07*10<sup>3</sup> (Ref. <sup>16</sup>) |  |
|  | *Ferrimonas kyonanensis* | WP_028114584.1 | 418 | Unclear (1) | N.D. | N.D. | N.D. | N.D. |
|  | *Shewanella putrefaciens* CN-32 | WP_011920089.1 | 414 | Unclear (1) | N.D. | N.D. | N.D. | N.D. |
| DddL | *Thioclava pacific* | WP_051692700.1 | 232 | $\geq 83$ (3) | N.D. | N.D. | N.D. | Mn<sup>2+</sup> |
| | *Rhodobacter sphaeroides* 2.4.1 | Q3J6L0.1 | 232 | $\geq 70$ (3) | N.D. | N.D. | N.D. | Mn<sup>2+</sup> |
| DddQ | *Ruegeria pomeroyi* DSS-3 | AAV94883.1 | 201 | 2~5*10<sup>-3</sup> (Refs. <sup>13, 18</sup>) | <20 (Ref. <sup>13, 18</sup>) | N.D. | N.D. | N.D. |
|  | *Ruegeria lacuscaerulensis* ITI-1157 | D0CY60 | 192 | 2.8*10<sup>-2</sup> (Ref. <sup>20</sup>) | 0.27 (Ref. <sup>20</sup>) | 39.1 (Ref. <sup>20</sup>) | 1.05*10<sup>-2</sup> (Ref. <sup>20</sup>) | Fe<sup>3+</sup> (Ref. <sup>20</sup>) |
| DddK | *Candidatus Pelagibacter ubique* HTCC1062 | WP_011281678.1 | 130 | ~30 (Ref. <sup>26</sup>) | 608 (Ref. <sup>26</sup>) | 5.1 (Ref. <sup>26</sup>) | 3.1 (Ref. <sup>26</sup>) | Ni<sup>2+</sup> (Ref. <sup>26</sup>) |
| DddW | *Ruegeria pomeroyi* DSS-3 | WP_011046214.1 | 152 | 61.2 Ref. <sup>25</sup> | 2.10*10<sup>3</sup> (Ref. <sup>25</sup>) | 4.5 (Ref. <sup>25</sup>) | 17.33 (Ref. <sup>25</sup>) | Mn<sup>2+</sup> (Ref. <sup>25</sup>) |

N.D. not determined
<sup>1</sup> Weak DMSP lyase activity was observed in crude lysates upon over-expression but activity was lost during purification. Attempts to reproduce the lysate activity gave ambiguous results.
<sup>2</sup> Ca<sup>2+</sup> is additionally required to obtain maximal activity, probably binding to a structural rather than the cupin site.
<sup>3</sup> Specific activity was estimated from the activity and protein concentrations in the crude lysate; see **Supplemental Fig. S3**.

Whilst the 1^st^ clade that includes AfDddY and DaDddY’s clearly encompasses highly active DMSP lyases, the identity of the 2^nd^ clade of putative DddY’s, including the *Shewanella, Ferrimonas* and *Synechococcus* genes, is unclear. Upon expression in *E. coli*, few of the representative genes we tested exhibited weak DMSP lyase activity in crude cell lysates (≤ 14 μM DMS min^1^mg^1^ total protein in crude lysate; compared to ~700 for DaDddY and ~400 for AfDddY). However, attempts to purify these enzymes resulted in very low yield and complete loss of activity. Replacing the signal peptide to PelB’s did not result in higher yield or activity. Attempts to optimize growth conditions for maximal crude lysate activity gave no improvement and even the initially observed activity failed to reproduce. Lack of activity could relate to misfolding, especially as the yield of soluble protein was low. Inferred ancestors were found to consistently yield proteins with high foldability and stability.^23^ We therefore inferred the ancestor of the *Shewanella* clade **(Figure 1A)** and expressed it in *E. coli*. However, the inferred *Shewanella* ancestor expressed poorly and was inactive similar to the *Shewanella putrefaciens* CN-32. Addition of various metal ions found to be essential for DaDddY (described below) to the growth medium or/and the lysis buffer had no effect either. Hence, at this stage it remains unclear whether the 2^nd^ clade is an integral part of the DddY family although its members are unable to correctly fold in *E. coli*. Alternatively, this clade represents a related enzyme family with some residual, promiscuous DMSP lyase activity. As described below, this ambiguity applies for other members of the Cupin-DLL superfamily that exhibit weak or no DMSP lyase activity upon expression in *E. coli*.

### DddY shares cupin motifs with other Ddd families

Having collected a range of DddY sequences, we could identify common sequence motifs, including the cupin motifs that are shared by DddQ, DddL, DddK and DddW^13, 25^ These cupin motifs could also be identified in DddY. Accordingly, these 5 ddd+ families share two conserved cupin-like sequence motifs **(Figure 2A).** As expected, the two remaining bacterial ddd+ families, dddD and dddP, and the algal Alma DMSP lyases, do not share these motifs as they belong to completely different superfamilies.

**Figure 2.**
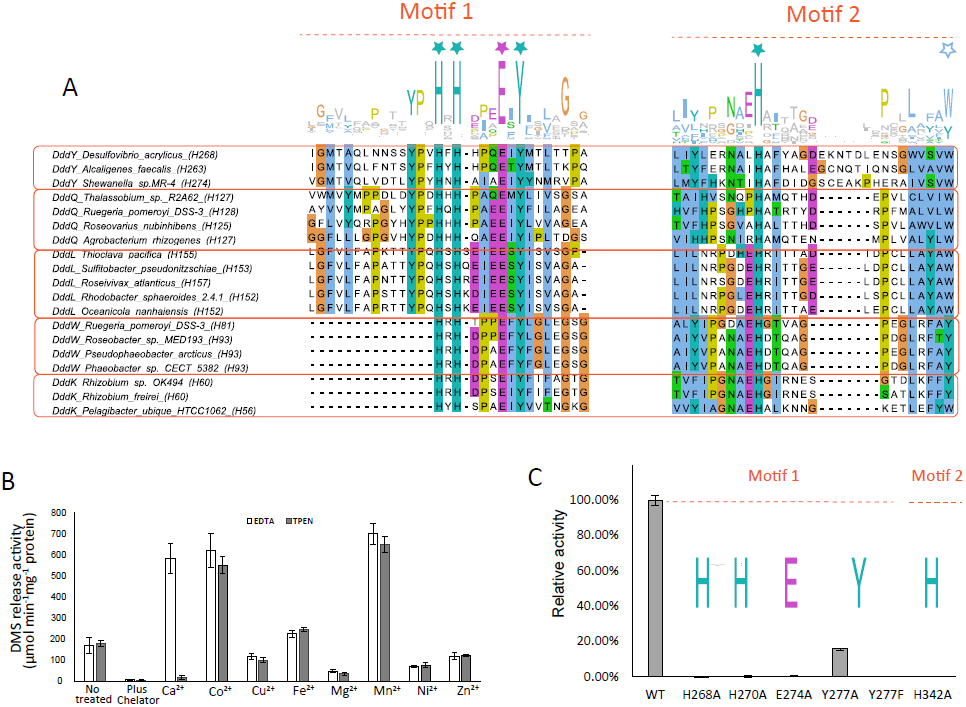
DddY is a cupin-like enzyme. (A) Sequence alignment of the two most conserved regions of the reported cupin Ddd+ families and DddY. The resulting sequence Logo is presented on top. Marked in asterisks are the 6 most conserved active-site residues. (B) Incubation of DaDddY with either EDTA or TPEN (2 mM) resulted in complete loss of activity. Addition of Ca^2+^ restored activity after EDTA treatment, whereas addition of transition metal ions restored activity after treatment with both chelators. (C) The activity of various mutants of DaDddY. Shown is relative activity compared to wild-type (μM DMS/min, at 100ng/mL enzyme; mutants were tested at 100ng/mL). Data points in panels B and C represent the average activity in 3 independent measurements and the error bars represent the S.D. values.

The first shared cupin motif comprises 4 entirely conserved residues – HxHxxxxExY (His268, His270, Glu274 and Tyr277 (numbering is for DaDddY). The second motif includes a third entirely conserved histidine, His342. Of the 5 cupin ddd+ families, structures are available for DddQ (PDB 4LA2, Ref.^19^; 5JSO and 5JSP, Ref.^20^; 4B29 and 5CU1), and DddK (PDB 5TG0, Ref.^26^). The structures indicate that two His residues, one from the 1^st^ motif and another from the 2^nd^ one, and the Glu of the 1^st^ motif, directly chelate the active-site metal ion (His141, Glu145 & His180 in PDB 5CU1). The second His of the 1^st^ motif and its Tyr (His270 and Tyr277 in DaDddY) are also within the active-site and in close proximity to the metal ion (< 4 Å, His125 and Tyr131 in PDB 4LA2; His141 and Tyr147 5CU1; His 150 and Tyr156 in PDB 4B29). Other conserved residues are part of both motifs. For example, a Trp/Tyr at the end of the 2^nd^ motif is also an active-site residue, with the Trp ring’s NH placed ~4.5Å from the metal ion (Trp363 in DaDddY; Trp178 in DddQ, PDB 4LA2; Trp195 in PDB 5CU1; Trp204 in PDB 4B29). The location of these residues within the active–site and close to the metal indicates that they are critical to function and also explains their absolute conservation in DddQ, DddL and DddY famillies.

Together, these two motifs, and foremost, the above-mentioned 6 active-site residues **(Figure 2A),** define the Cupin-DLL superfamily. Further analysis of this superfamily’s gene content beyond the 5 Ddd+ families is provided in the last section of the Results.

### DddY demands a transition-metal ion cofactor

Given DddY’s sequence assignment as a cupin DMSP lyase, its catalytic activity should be utterly dependent on a transition metal ion.^27^ Indeed, following incubation with the metal chelator EDTA, DaDddY lost all DMSP lyase activity **(Figure 2B).** However, confusingly, enzymatic activity was restored, and even augmented relative to the untreated enzyme, not only by transition metals (e.g. Co^2+,^ Fe^2+,^ Mn^2+)^ but also by Ca^2+^ that was added as a negative control. It is unlikely that a cupin metal site will bind or be fully active with both an alkali and a transition metal ion. We thus surmised that Ca^+2^ binds at an alternative site that is permissive to both alkali and transition metal ions, while EDTA actually failed to remove the transition metal ion at the cupin site. Indeed, when TPEN, a chelator selective for transition metals, was applied, the DMSP lyase was completely lost and could be fully recovered by addition of transition metal ions, but not by Ca^2+^ **(Figure 2B).** This result confirmed that DddY’s activity is dependent on a transition metal cofactor, as observed with all cupin-like enzymes, including DddW and DddQ.^13, 19^ The alternative metal binding site is likely required for maintaining DddY’s structure, and may be unique to this family whose size is far larger than any other Cupin ddd+ **(Table 1).**

### The conserved motifs are essential for the DMSP cleavage activity

To confirm the role of the most conserved amino acids in the active site of DddY, a series of site-directed mutants to Ala was generated at DaDddY’s background. The mutations H268A, H270A, E274A and H342A all showed complete loss of the activity, as reported for other cupin Ddd+ enzymes such as DddW.^25^ The Y277A mutation retained around 20% of the wild-type’s activity. However, when Tyr277 was replaced by phenylalanine whose side-chain is similar to tyrosine, enzymatic activity was completely lost **(Figure 2C).** The CD spectra of all these mutants were essentially identical to wild-type, suggesting that the mutations perturbed the active-site rather than the enzyme’s overall fold **(Supplemental Fig. S2A).** These residues of the two cupin motifs are therefore essential for enzymatic activity, foremost, for the ligation of the active-site metal ion.^27^ However, Tyr277 that belongs to the first motif presents a puzzle. This tyrosine has been suggested to play a key role in proper positioning of the transition metal ion and the substrate in DddQ’s active-site (Tyr131 is PDB 4LA2, 5JSO and 5JSP; Ref. ^19^,^20^).^20^ However, its substitution to Ala in DaDddY has only a minor effect. Unexpectedly, however, the more conservative substitution to Phe led to complete inactivation, possibly because in the absence of the interacting hydroxyl of Tyr, a hydrophobic moiety in the metal’s close vicinity is highly deleterious. To get a clear understanding of the role of this active-site tyrosine, its mutants in DddK, DddW and DddL were examined. Similarly to DddY, the Tyr277 to Ala mutants showed a relatively mild decrease in activity around 10% residual activity). In DddL and DddW, the Phe mutants showed near-complete loss of activity as in DaDddY, whereas in DddK replacements to either Ala or Phe had similar effect (around 10% residual activity, **see Supplemental Fig. S2B).** Overall, it appears that the Tyr277 of the 1^st^ motif plays a role in DaDddY’s DMSP lyase activity, as well is in the other Cupin Ddd+ enzymes, but its role is secondary, certainly in comparison to the metal-ligating residues (His342 in DaDddY, His163 in DddQ, PDB 4LA2).

### DddL is potentially a highly active DMSP lyase

The shared cupin motifs, and the role and location of their conserved residues, indicate that DddY, DddQ, DddL, DddK, and DddW belong to the same superfamily. However, as shown for a number of enzyme superfamilies, common evolutionary origin typically results in overlapping activities whereby the native function of one family is observed as promiscuous in related one(s), and vice versa. ^28-31^ The levels of DMSP lyase activity in Cupin-DLL family members range over 6 orders-of magnitude **(Table 1).** While enzymatic parameters are widely distributed, the *kcat*/*KM* values within enzyme classes do not vary as widely; foremost, *kcat*/*KM* values of < 1 M^-1^s^-1^ are highly unlikely to reflect the native substrate (the average *kcat*/*KM* value is ~ 10^5^ M^-1^s^-1.32^ This suggest that whereas DddY’s native function is almost certainly as DMSP lyase (*kcat*/*KM* ≈ ^10^^6^ M^-1^s^-1^), DddQ’s DMSP lyase activity is likely to be promiscuous (*kcat*/*KM* < 1 M^-1^s^-1^; Table 1). To complete the picture with respect to activity levels, we tested the DMSP lyase activity of members from all five Ddd+ families. Foremost, we sought to determine DddL’s activity,^32^ as to our knowledge, reports of its specific activity, let alone of its kinetic parameters, are not available.

We examined two DddL orthologues: the previously described *Rhodobacter sphaeroides* DddL (RsDddL)^11^, a DddL gene we identified in the *Thioclava pacifica* genome (TpDddL; 78% amino acid identity to *R*. *sphaeroides* DddL), and *Sulfitobacter* EE-36 DddL (50% amino acid to RsDddL).^11^ RsDddL was cloned and expressed in *E. coli* with His-tags at either the N- or C-terminus, or an N-terminal Strep-tag. High DMSP lyase activity was repetitively observed in crude cell lysates, indicating a soluble, correctly folded and active enzyme. However, all attempts to purify RsDddL failed. The C-terminal His-tag construct did not bind to Ni-NTA beads, the N-terminal His-tag construct was insoluble and inactive, while the N-terminal Strep-tag construct bound the resin, but its enzymatic activity was lost upon elution **(Supplemental Fig S3).** Subsequently, TpDddL and *Sulfitobacter* DddL was cloned and expressed with an N-terminal Strep-tag, but showed similar behavior. Various additives were tested, including detergents and metal ions, but enzymatic activity could not be retained during purification. Nonetheless, the specific activity of both DddL’s in freshly prepared *E. coli* crude cell lysates, as estimated from its protein concentration and DMSP lyase activity, seems high, in the range of 70 Units **(Supplemental Fig S3).** The kinetics indicated a linear increase of initial rates with DMSP concentrations up to 10 mM suggesting a high *K*_*M*_ and accordingly high *k*_*cat*_. This value of specific activity is probably underestimated, as the soluble fraction is likely to contain misfolded enzyme. Nonetheless, this specific activity is up to 10-fold higher than DddK and DddW’s, and is well above 1000-fold higher than DddQ’s **(Table 1).**

### Genome context suggests that DddQ and DddW’s primary activity is not DMSP lyase

In bacterial genomes, enzymes of the same pathway are typically found in gene clusters or even in the same operon. Thus, genome context can provide important hints regarding the enzymatic function of *ddd+* genes. Acrylate, the verified product of all cupin Ddd+ enzymes **(Table 1),** is toxic,^33, 34^ and previous studies indicated proximal genes that relate to acrylate catabolism. These include: *acuI*, a zinc/iron dependent alcohol reductase that converts acrylyl-CoA into propanoyl-CoA;^17, 33^ acuK (a dehydratase) and *acuN* (a CoA transferase) that can jointly convert acrylate into 3-hydroxypropionate;^35^ or cytochrome dependence oxi-reductases that could also catabolize acrylate.^36^

We applied few common tools of genome context analysis,including EFI-GNT,^37^ STRINGE,^38^ EASYFIG,^39^ and RODEO.^40^ Of these, RODEO provided the most systematic results for the examined Ddd+ genes, including the previously studied DMSP CoA-transferase/lyase *dddD* that serves as benchmark for a *bona fide* DMSP lyase.^9^ Accordingly, *dddD* is characterized by proximal *dddB* and *dddc* genes, encoding an iron-containing dehydrogenase and a methyl-malonate semi-aldehyde dehydrogenase-like protein, respectively **(Supplemental Fig S5A;** Refs.^2, 7^). The product of DddD, 3-hydroxypropionate-CoA, is converted into malonate semi-aldehyde (DddB) and then to acetyl-CoA (DddC).^2^ Accordingly, *dddY*’s genome neighborhoods repetitively contain putative acrylate utilizing genes (Ref ^17^; **Supplemental Fig S5B).** Indeed, the bacterium in which DaDddY originally resides (*Desulfovibrio acrylicus*) converts acrylate into propionate;^41^ and *Alcaligenes faecalis* M3A strain that contains AfDddY converts acrylate into 3-hydroxypropionate.^42^

In DddL’s case, however, the genome context is not conserved, making it hard to draw a clear conclusion (in 4 out of 15 genomes an *acuI*-like zinc containing reductase is present just next to DddL; **Supplemental Fig. S6A).** Similarly, because DddK sequences are only found in several strains of *Pelagibacter ubique* (SAR11),^12^ it is difficult to get a systematic prediction based on genome context. In the genome of SAR11, the proximity ofenoyl-ACP-reductate,β-ketoacyl-ACP-synthase and β-hydroxydecanoyl-ACP dehydratase indicate a relation to fatty acid or polyketide biosynthesis. ^43^ **(Supplemental Fig. S6B)**

The genome neighborhoods of both *dddQ* and *dddW* could be consistently derived. For DddW, the proximal D-alanyl-D-alanine carboxypeptidase gene **(Supplemental Fig. S6C)** suggests a gene cluster that is involved in bacterial peptidoglycan biosynthesis. The conserved neighborhood of dddQ, on the other hand, includes a putative mandelate racemase-like protein and a putative dimethylglycine dehydrogenase **(Figure 3).** The mandelate racemase-like gene exhibits present distinct homology (40% and 35% amino acid identities, respectively) to two recently identified enzymes: cis-3-hydroxy-L-proline dehydratase and 4-hydroxyproline betaine 2-epimerase.^44, 45^ These two neighbors of *dddQ*, and the dimethylglycine dehydrogenase-like neighbor, suggest that DddQ takes part in the catabolism of proline-betaine or/and hydroxyproline-betaine. Its DMSP lyase activity is therefore likely to be promiscuous, as also indicated by its markedly low activity **(Table 1)** and lack of selectivity (see accompanying paper).

**Figure 3.**
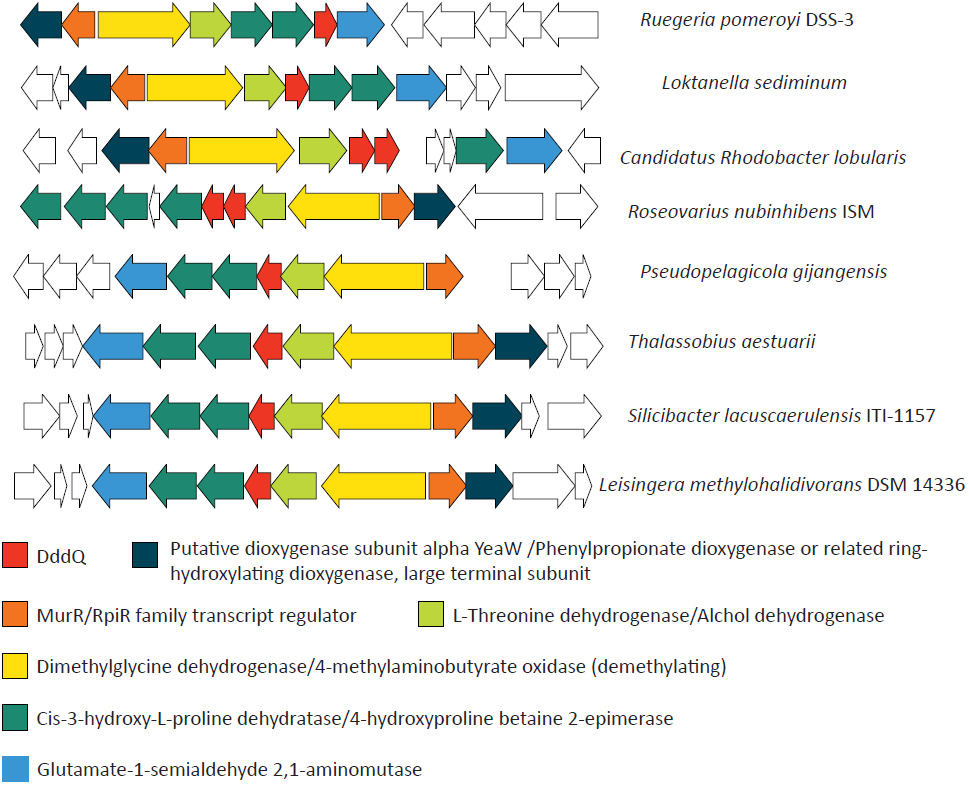
Our search identified 8 dddQ-like genes in sequenced genomes. Genome context analysis was performed by RODEO^40^, annotation of each genes were obtain from the original genome annotation, and manually checked. The genes encoding orthologues are highlighted with the same colour. The result suggests that *dddQ* gene is associated with the catabolism of proline-betaine or/and hydroxyproline-betaine. Shown are the results.

### Other putative members of the Cupin-DLL superfamily

To gain further insights into the evolutionary relationships of the cupin DMSP lyases, pfam model PF16867 was used to search against the NCBI protein database and the Tara marine metagenomics database.^46^ Following filtering, the remaining sequences were aligned (361 sequences in total) and an extended profile that encompassed all potential superfamily members was generated (see **Methods** and **Supplemental Dataset 2).** A phylogenetic tree was also built **(Supplemental Fig. S4).** Needless to say that a tree that includes sequences with as low as 15% identity and highly variable lengths (90 - 441 amino acids) is inaccurate. Nonetheless, it provided an initial view of the content of this superfamily and of where the Ddd+ families map within it, as schematically summarized in **Figure 4.**

**Figure 4.**
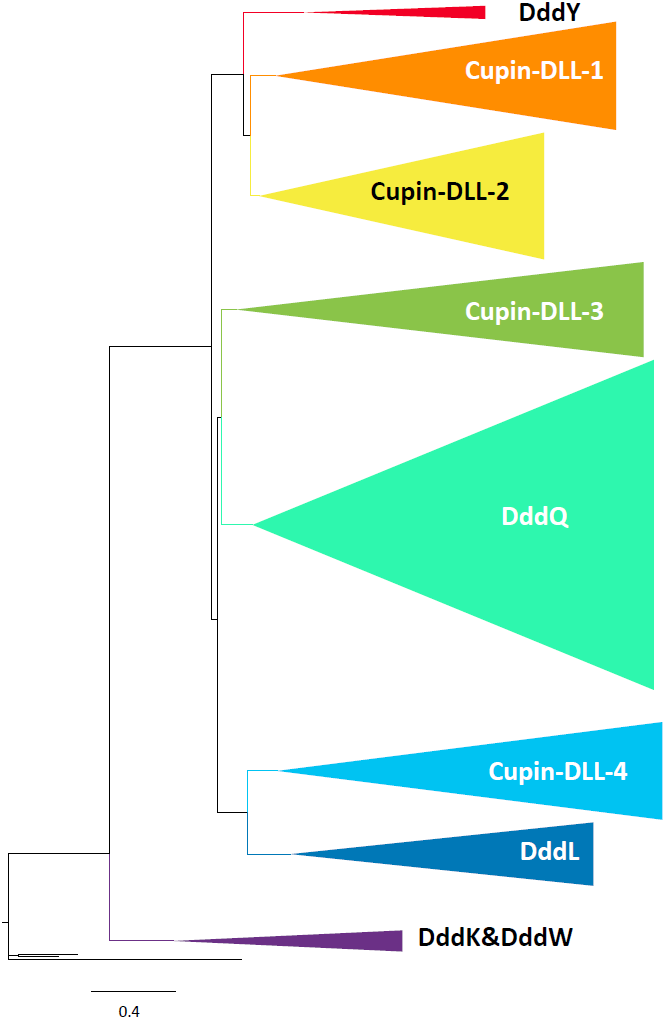
A schematic phylogenetic tree of the Cupin-DLL family (based on the phylogentic tree presented in **Supplemental Figure S4).** Overall, 8 clades could be identified that include the 5 known Ddd+ families, whereby DddK and DddW belong to the same clade. Of the 4 Ddd+ clades, only DddY and DddL seem to be specialized DMSP lyases. The primary function of the other Ddd+ clades, and foremost of DddQ, and of the 4 additional clades (marked arbitrarily as 1-4), remains unknown.

**Figure 5.**
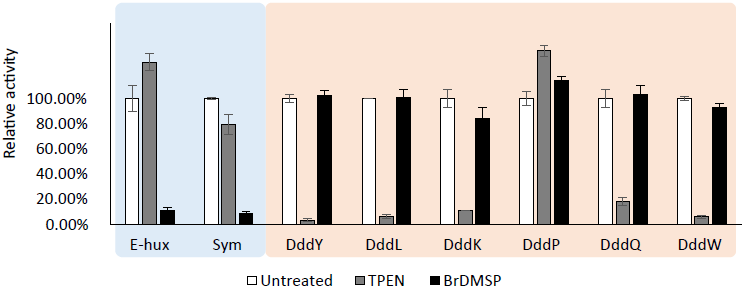
Cupin Ddd+ DMSP lyases are selectively inhibited by the metal chelator TPEN. All tested enzyme were incubated with 10uM Br-DMSP or 1mM TPEN (100mM Tris, 100mM NaCl, 1 mM CaCl2, pH 8.0). The residual DMS release rates were tested after 1 hour incubation at ambient temperature. Shown is the relative DMSP lyase activity compared to the untreated enzyme. The Alma DMSP lyases from *Emiliania huxleyi* and *Symbiodinium* are marked as E-hux and sym, respectively.

The cupin-DLL superfamily is large, and its various clades are widely diverged, suggesting high functional diversity. DddQ is by far the largest family. With the exception of DddW and DddK that are closely related, the other 4 Ddd+ families are sporadically distributed throughout the tree, suggesting an independent emergence of DMSP lyases from enzymes with another function, at least for DddL and DddY. However, the vast majority of Cupin-DLLs are obviously not DMSP lyases. However, as exemplified by DddQ, the shared evolutionary origin can give rise to promiscuous DMSP lyase activity.

### Cupin-DLL enzymes are inhibited by the metal chelator TPEN and unaffected by Br-DMSP

We have recently reported that 2-bromo-3-(dimethylsulfonio)-propionate (Br-DMSP) is a potent and selective mechanism-based inhibitor of the algal Alma DMSP lyases, and does not inhibit any known bacterial Ddd+ enzymes.^6^ This inhibitor, that covalently modifies the active-site cysteine of the Alma lyases, is a biochemical probe that could detect the presence of Alma DMSP lyase in various marine organisms and environments and assess their contribution to DMS release. We sought to verify that TPEN **(Figure 2B)** is a complementary probe that could identify the cupin DMSP lyases. To this end, DddY, DddL, DddQ, DddK and DddW were incubated with either Br-DMSP or with the transition metal chelator, TPEN. The reaction buffer also contained CaCl2 to avoid Alma or DddY losing activity (likely due to a structural calcium site; see **Figure 2B).** The Alma DMSP lyases from *Emiliania huxleyi^8^* and *Symbiodiniome*^6^ were examined as control. All cupin Ddd+ enzymes were all inhibited by TPEN while Br-DMSP had no effect. Conversely, both Alma DMSP lyases were inhibited by Br-DMSP and unaffected by TPEN. Surprisingly, DddP, as a member of the metallo-enzyme superfamily M24 peptidase (see also accompanying paper), showed no inhibition by TPEN; a similar result has been reported in Refs.^47^,^48^, neither EDTA nor 1,10-phenanthroline inhibited DddP’s activity. The selective pattern of inhibition by Br-DMSP and TPEN can therefore be applied to identify the presence of the two major classes of DMSP lyases, Alma and cupin-DLL, in different marine environments and species, be they algae or bacteria. This biochemical profiling can complement genomic data to unravel the origins of marine DMS release.

### Discussion

The search for DddY’s affiliation led to the identification of an enzyme superfamily dubbed Cupin-DLL **(Figure 4).** With the exception of DddD and DddP, all bacterial ddd+ genes implicated as DMSP lyases belong to this newly identified superfamily. We have also identified TPEN as a common inhibitor of the cupin Ddd+ enzymes.

The phylogenetics suggest that DMSP lyases evolved within this superfamily independently along multiple lineages. This phenomenon of parallel evolution is commonly observed in enzyme superfamilies.^30, 49^ Its origins are in shared promiscuous activities - relatively distant superfamily members often share the same promiscuous activity. Such promiscuous activities comprise the starting point for the divergence of new enzymes by turning a latent, promiscuous activity into a physiologically relevant function, initially alongside the enzyme’s original function. Indeed, bi-functional, ‘generalist’ enzymes commonly comprise evolutionary intermediates, although they may persist for long periods as is apparent by the dominance of multifunctional enzymes in extant proteomes.^28^ As the divergence process proceeds, highly active, selective ‘specialist’ enzymes evolve, typically by duplication and sub-functionalization of the bi-functional generalist ancestor^50^ Parallel evolution, as well as convergent evolution as exemplified by the independent evolutionary origins of the algal Alma1 DMSP lyase, also relates to DMSP lysis being a facile reaction,^2^ and foremost to the fact that lysis is initiated by abstraction of a proton from a carbon next to a carboxylate (a-carbon). This is one of the most common steps in enzyme catalysis, and several large enzyme superfamilies share it as the key catalytic step, including the Asp/Glu racemase superfamily to which the Alma DMSP lyases belong. Accordingly, DMSP lyases that belong to the enolase superfamily – whereα-proton abstraction is the hallmark, are also highly likely to exist. ^51^

Further, our view as evolutionary biochemists is that the Ddd+ families likely represent an entire range of evolutionary states – they may be fully diverged DMSP lyase specialists (DddY and DddL), bi-functional intermediates (possibly DddK/W) or enzymes with a different function that only exhibit latent, promiscuous DMSP lyase activity (DddQ in most likelihood). The level of activity is an important parameter with respect to physiological relevance. However, low specific activity may also be due to suboptimal folding or/and reaction conditions. We thus pursued an independent test of substrate profiling. As described in the accompany paper, it appears that activity levels and substrate selectivity are correlated – the least active enzymes, foremost DddQ, also show the weakest signature of a DMSP tailored active-site.

DddK and DddW reside in the mid-range by both criteria, activity and selectivity, and could therefore be bi-functional enzymes. Enzymes in secondary metabolism tend to exhibit lower catalytic efficiency compared to central metabolism enzymes^32^ and also exhibit multi-functionality. Indeed, while their DMSP lyase activity of DddK/W is modest (*kcat*/*KM* ≈ 10^3^ M^-1^s^-1^), their rates are notably comparable to the rates of DMSP demethylation by the corresponding enzyme (DmdA).^26^ This comparison is relevant because DMSP is catabolized via two routes, lysis to give DMS as described here, or demethylation to give methylmercapto propionate,^2^,^4^, ^52^ and the demethylation route is actually the dominating one.^2^, ^3^ ^53^

While following the identification of the cupin-DLL superfamiliy the picture with respect to the DMSP lyases has become somewhat clearer. However, the key remaining question is the primary function of the cupin-DLL superfamily, and specifically of dddQ that also comprises its largest clade **(Figure 4)**. Future research may reveal the function of Cupin-DLLs that are not DMSP-lyases, and may thus also shade light on the evolutionary history and function of the ddd+ families.

## Material and Methods

### Phylogenetic analysis

Hypothetical cupin-like DMSP lyases sequence were collected by using Hidden Markov Model based search using the HMMsearch of the HMMer package.^54^ Briefly, the existing HMM model of DMSP lyase, PF16867, was used to search the NCBI non-redundant protein database and the Tara metagenomics database.^46^ All hits were collected, aligned, and manually filtered by the presence of the two conserved motifs and minimal length (≥ 90 amino acids). The sequence redundancy in the remaining set was minimized using CD-hit with 70% cutoff. The resulting 361 sequences were aligned by MUSCLE,^55^ followed by minimization of gaps (manual removal of sporadic insertions) to obtain a core alignment of 115 amino acids length (the smallest known Cupin Ddd+ enzymes (DddK’s) are 130 amino acids length). A phylogenetic tree were built from this core alignment using the Markov chain Monte Carlo (MCMC) method of the MrBayes program.^56^

### Enzyme cloning and mutagenesis

All tested DddY genes (including D*adddY*, Af*dddY*, *Shewanella dddY*, *Ferrimonas dddY*, and *Synechococcus dddY*) were synthesized by Gen9. The *Ruegeria pomeroyi* DSS-3 *dddQ* and *Rhodobacter sphaeroides* 2.4.1 *dddL* genes were kindly provided by Professor Andrew Johnston, University of East Anglia. The synthesized fragments were amplified and the PCR product was digested with *Nco*I and *Hind*III, and then cloned into the expression vector pET28a for expression with C-terminal His tag. These plasmids were also used as the template for site-directed mutagenesis. The *TpdddL* was synthesized by Gen9 and contained an N-terminal Strep-tag and a stop codon before the *Hind*III site. It was cloned into pET28a via the *Nco*I and *Hind*III sites. Point mutations in *dddY*, *TpdddL*, *dddK* and *dddW* were introduced with mutagenesis oligos using SOE-PCR-based approach. The PCR fragments were digested with *Nco*I and *Hind*III and cloned into pET28a. All mutants were verified by DNA sequencing.

### Enzyme purification

Enzymes were typically expressed using pET28a plasmid in *E.Coli* BL21 (DE3). Cells were grown for overnight in 5 mL LB medium at 37°C. These cultures (1 mL) were used to inoculate 1 liter LB cultures that were subsequently grown at 37°C to OD_600nm_ of 0.6-0.8. The growth temperature was reduced to 16°C, and enzyme expression was induced with 0.1mM IPTG. Following overnight growth at 16°C, the cells were harvested by centrifugation at 4° C. DddY were purified on Ni-NTA beads. Briefly, cells were re-suspended in 50mL lysis buffer (100mM Tris-HCl pH 8.0, 100mM NaCl, 1mM CaCl_^2^_, 10 mg lysozyme and 10 μg bezonase). Cell suspensions were incubated in ambient temperature for 30 min and sonicated. Lysates were clarified by centrifugation and loaded on 2 mL Ni-NTA agarose beads (Millipore). Binding was performed at 4°C, beads were washed with 50 mL lysis buffer followed by 100 mL lysis buffer with 35 mM imidazole. The enzyme was eluted with 150 mM imidazole. Fractions were analyzed by SDS-PAGE and activity, combined, and the purified enzyme was concentrated by ultrafiltration (Amicon). For TpDddL with N-terminal Strep-tag, clarified cell lysates were loaded on 2mL Strep-tactin beads (IBA), and the beads were rinsed with 50mL lysis buffer. The protein was eluted with 2.5mM desthiobiotin. Final enzyme concentrations were determined by the BCA assay.

### DMSP lyase activity assays

DMS release was measured as previously described^10^ Briefly, freshly prepared 100 mM Tris-HCl pH 8.0 with 100 mM NaCl was used for the enzymatic assays supplemented with 10 mM DMSP as default. The high buffer capacity is critical for this assay, because DMSP as applied as hydrochloride salt, and its cleavage releases protons. Reactions were performed at 30°C (typically for 5 min,) and terminated by 1000-fold dilution into 30 mL chilled 10mM glycine pH 3.0 in sealed glass vials. Enzyme concentration were typically as follows: *Ehux*-Alma1, 0.3ug/mL; *Sym*-Alma1, 1μg/mL; *Desufovibrio* DddY 20ng/mL; *Alcaligenes* DddY 50ng/mL; DddW, 15μg/mL; DddQ, 100μg/mL; DddK, 8μg/mL. DddL was assayed in crude lysate, at an estimated concentration of 8ug/mL **(Supplemental Fig. S3).** DMS levels were determined using an Eclipse 4660 Purge-and-Trap Sample Concentrator system (OI Analytical) followed by separation and detection using GC-FPD (HP 5890) equipped with RT-XL sulfur column (Restek). All measurements were calibrated using DMS standards.

### Metal chelation and complementation

Protein samples were incubated with 1mM EDTA or TPEN, at 30 °C for 1 hr. The chelators were removed by dialysis at 4 °C overnight. Different metal ions were supplemented by adding the corresponding chloride salts to the dialyzed apo-proteins at 2 mM concentration, and incubating at 30 °C for 1 hr. DMSP lyase activity was subsequently measured.

